# Variability in cognitive task performance in early adolescence is associated with stronger between-network anticorrelation and future attention problems

**DOI:** 10.1101/2022.07.07.499196

**Authors:** Sarah E. Chang, Agatha Lenartowicz, Gerhard S. Hellemann, Lucina Q. Uddin, Carrie E. Bearden

## Abstract

**Background:** Intra-individual variability (IIV) during cognitive task performance is a key behavioral index of attention and consistent marker of ADHD. In adults, lower IIV has been associated with anticorrelation between the default mode network (DMN) and dorsal attention network (DAN) - thought to underlie effective allocation of attention. However, whether these behavioral and neural markers of attention are (i) associated with each other and (ii) can predict future attention-related deficits has not been examined in a developmental, population-based cohort.

**Methods:** We examined relationships at the baseline visit between IIV on three cognitive tasks, DMN-DAN anticorrelation, and parent-reported attention problems using the Adolescent Brain and Cognitive Development Study (n=11,878 participants, aged 9-10, female=47.8%). We also investigated whether behavioral and neural markers of attention at baseline predicted attention problems, 1, 2, and 3 years later.

**Results:** At baseline, greater DMN-DAN anticorrelation was associated with lower IIV across all three cognitive tasks (unstandardized β: 0.22-0.25). Older age at baseline was associated with stronger DMN-DAN anticorrelation and lower IIV (β: -0.005--0.0004). Weaker DMN-DAN anticorrelation and IIV were cross-sectionally associated with attention problems (β: 1.41-7.63). Longitudinally, lower IIV at baseline was associated with less severe attention problems, 1-3 years later, after accounting for baseline attention problems (β: 0.288-0.77).

**Conclusions:** The results suggest that IIV in early adolescence is associated with worsening attention problems in a representative cohort of US youth. Attention deficits in early adolescence may be important for understanding and predicting future cognitive and clinical outcomes.

## Introduction

Momentary lapses in attention can interfere with goal-directed behaviors. In individuals with attention deficits, these lapses are persistent and can hinder task completion at work or concentrating in school. A key metric of attentional lapses is intra-individual variability (IIV), which is defined as trial-by-trial fluctuation in reaction time during a timed cognitive task (1). Higher IIV is a robust behavioral marker of attention-deficit/hyperactivity disorder (ADHD), ubiquitous across types of speeded reaction time tasks, across contexts and environments (Hedge’s *g* = 0.76; ADHD vs. controls) (2,3). Importantly, a meta-analysis of 319 studies indicates that higher IIV in individuals with ADHD is not driven by mean reaction time (3). Across the lifespan, IIV exhibits a U-shaped trajectory, with dramatic reductions during childhood and adolescence corresponding with a major developmental shift in sustained attention (4-7). At the same time, adolescence is a sensitive period involving increased risk for onset of many psychiatric disorders (8). Investigating the spectrum of attention variability at this juncture of development may be important for understanding and predicting future cognitive and clinical outcomes.

While greater IIV has been established as a robust marker of attention deficits, its neural bases remain an open question. Studies using resting state functional magnetic resonance imaging (rsfMRI) implicate dysfunction of functional brain networks in greater IIV and attention problems (9-14). In particular, rsfMRI studies have identified the default mode network (DMN) and task-positive networks, including the dorsal attention network (DAN) and the balance between these networks as key for attention allocation (11). The DMN is engaged during internally-directed processes, like self-referential processing and the recollection of previous experiences, and its activity is typically reduced when individuals focus on tasks requiring visuospatial attention (15-18). The DMN consists of the mPFC, posterior cingulate cortex, temporoparietal junction, lateral and medial temporal lobes (15, 17, 18). The DAN consists of the frontal eye fields and the inferior parietal sulcus and is involved in external, top-down visual attention (19-21).

The DMN and DAN typically exhibit negatively correlated (‘anticorrelated’) patterns of activity: as activity in the regions of the DAN increases when engaging in an attention-demanding task, activity in the DMN decreases (17, 22-25). Evidence from fMRI and EEG studies suggest that excess DMN activity during tasks is associated with less efficient cognitive performance (26-29). Furthermore, DMN deactivation during attentional tasks is associated with better cognitive performance (30,31). The anticorrelation of these two networks may be a fundamental property of brain organization that supports effective attention allocation, by focusing on the task at hand and suppressing internally-directed thoughts to support neurocognitive performance (11, 17, 32).

Consistent evidence from rsfMRI research across the lifespan and studies of ADHD links DMN-DAN anticorrelation with attention allocation. For instance, these networks become increasingly anticorrelated in infancy through adolescence and less anticorrelated through older adulthood (33-35). Kelly et al. 2008 was the first to link DMN-DAN anticorrelation at rest with reduced IIV in healthy adults. This finding has been replicated and extended through task-based fMRI and EEG studies (34, 36-40). Recently, Owens et al. 2020 linked DMN-DAN anticorrelation with less severe parent-reported attention problems using the ABCD study at baseline. From the ADHD literature, a meta-analysis found that anticorrelation between DMN and task-positive networks was diminished in children and adolescents with ADHD compared with typically developing (TD) youth (42). Furthermore, evidence from dynamic functional connectivity (FC) analyses, a method that measures moment-to-moment shifts in connectivity (43), suggests that children with ADHD spent less total time in, and switched out of, anticorrelated states involving the DMN and task-relevant networks more frequently, compared with TD children (44). Lastly, preliminary evidence from 11 drug naive children with ADHD and 11 TD children indicates that methylphenidate (a stimulant used to treat ADHD) initiated anticorrelation between DMN and task positive networks, which appeared to reduce intra-individual variability (45).

While this cogent body of literature supports the notion that DMN-DAN anticorrelation is linked with IIV, several clinically relevant questions and literature gaps remain. While IIV is a robust marker of categorical ADHD diagnosis (ADHD vs. TD controls), it’s unclear whether IIV can be used in the general population to predict future attention problems (3). Case-control study design does not capture the full spectrum of attention problems because individuals with milder symptoms that do not reach clinical impairment may be excluded (40, 44-45). Importantly, studies linking mean DMN-DAN anticorrelation at rest to IIV have only been established in adulthood and are relatively small (11, 37). Yet, studying IIV in early adolescence is particularly important. Evidence from executive function task performance from n>10,000 participants, aged 8-35, has shown that executive function (including inhibitory attention) undergoes the most rapid development in early adolescence (46). Whether DMN-DAN anticorrelation develops alongside, prior to, or in response to this dynamic development is unknown. Lastly, whether DMN-DAN anticorrelation can be used to predict future attention problems has not been examined in a large, population-based sample. Deviations in these neural or behavioral markers of attention during this important stage of cognitive development may lead to poorer cognitive outcomes in adulthood.

To address these questions, we examine the relationship between intra-individual variability (IIV) in timed cognitive tasks and resting-state FC from the ABCD study, a population-based, longitudinal cohort of 11,878 9-10 year olds. In contrast to previous work, this study captures the full spectrum of attention problems during a period of dynamic cognitive and neural development (47, 48). The main hypotheses using cross-sectional data at baseline were the following: (1a) Stronger DMN-DAN anti-correlation will be associated with lower IIV, across cognitive tasks; (1b) within these models, older baseline age will be associated with stronger DMN-DAN anticorrelation and (1c) lower IIV. Secondly, we examined the cross-sectional relationship of lab-based cognitive and neuroimaging measures (IIV and DMN-DAN anticorrelation) with parent-report of their child’s attention problems. We predicted that both (2a) weaker DMN-DAN anticorrelation and (2b) higher IIV would be associated with more severe attention problems. Lastly, we tested two hypotheses using longitudinal data: (3a) weaker DMN-DAN anticorrelation and/or (3b) greater IIV would be associated with future attention symptoms, at 1-year, 2-year and 3-year follow-up timepoints, after accounting for baseline attention symptoms. In an exploratory analysis, we also examined whether IIV and/or DMN-DAN anticorrelation may be linked with externalizing symptoms, as deficits in attention may contribute to problems with self-regulation more broadly (49-51). Finally, when investigating behavioral and neural measures of attentional variability, we examined other key factors associated with these variables, including sex, race/ethnicity and parental income, to better contextualize these associations.

## Methods

### Participants

Data from 11,878 participants were obtained through the ABCD study, a prospective, longitudinal study tracking 9-11 year olds for the following 10 years, across 20 research collection sites in the United States (52). Each study site obtained informed consent from parents and assent from children, approved by each site’s Institutional Review Board (IRB), with centralized IRB approval at the University of California, San Diego. All de-identified raw and processed data for these analyses were from ABCD Data Release 4.0, accessed through the National Institute of Mental Health Data Archive (Collection 2573). Due to accessing and analyzing de-identified data, this study was exempt from IRB approval. Exclusion criteria for these analyses included: (1) missing or not reported demographics including race/ethnicity information (n=2) or parental income (n=1022); (2) missing trial-wise neuropsychological test data for the Flanker (n=1310), Pattern Comparison Processing Speed (n=1316), Dimensional Change Card Sort (n=1687); or (3) missing psychiatric symptom data at baseline (n =10). In concordance with previous work, additional subjects were removed from imaging analyses if they (1) had more than 600 frames of their resting-state scans with framewise displacement (FD) of 0.2 mm or higher, as previously published (n=202, 53-54), or (2) were scanned on a Philips scanner due to post-processing errors, as recommended by the ABCD Data Analysis and Informatics Core (DAIC) (n=1938, NDA Release Notes for Data Release 4.0, 55). While the ABCD study offers unprecedented sample size with FC summary metrics, neurocognitive battery, and psychiatric assessments, we constrained these analyses to variables most relevant to the investigation of IIV to avoid statistical problems with exploratory approaches. Baseline demographic characteristics are reported in Table 1; see the Supplement for study demographic information for year 1, 2, and 3 follow-up timepoints.

**Table 1.**
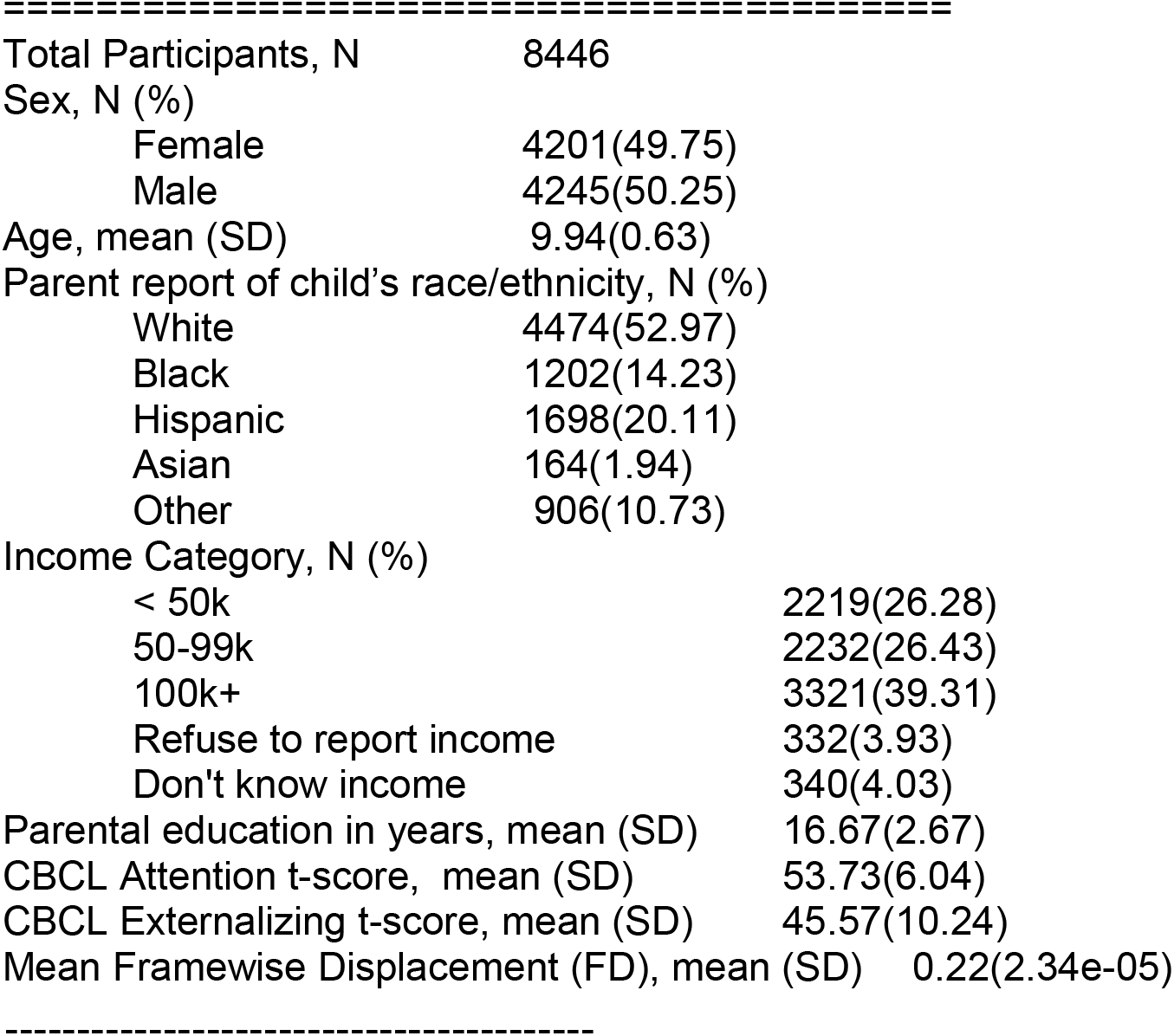
Baseline Demographics Table, based on imaging inclusion criteria.

### Measures

#### Neuropsychological Test Battery

Participants completed a developmentally-appropriate neurocognitive battery, the National Institute of Health (NIH) Toolbox (52). The NIH Toolbox cognitive battery includes 7 tasks, covering episodic memory, executive function, attention, working memory, processing speed and language abilities. This neurocognitive battery was designed to comprehensively assess these domains, has been used in longitudinal studies across childhood and adolescence, and is psychometrically sound (52).

The current study used the timed-reaction time tasks from NIH Toolbox cognitive battery, which includes the Flanker, Pattern Comparison Processing Speed, and Dimensional Change Card Sort tasks. The Flanker and Dimensional Change Card Sort task show excellent test-retest reliability (ICC=0.92 for both measures) and the Processing Speed task shows good test-retest reliability (ICC=0.84) in children and adolescents over a two-week interval (56, 57). Details of these tasks have been published elsewhere (52, 58).

Intra-individual variability for each task and participant was operationalized as the standard deviation in reaction time across all correct trials. To remove extremely large outliers, IIV was winsorized at three standard deviations for each task. Because the Flanker task assesses attention specifically, we focused first on testing our analyses using Flanker IIV. Then, we examined associations using IIV from the Dimensional Change Card Sort and the Pattern Comparison Processing Speed tasks, to assess whether the associations would generalize to these related cognitive domains.

#### Imaging Procedure: Acquisition

ABCD imaging collection, acquisition, and analysis has been previously described (59-61). All participants were scanned on 3T scanners, including Prisma (Siemens), Discovery MR750 (General Electric), Achieva dStream or Ingenia CX (Philips) with a 32-channel head coil. Participants completed T1-weighted and T2-weighted structural scans, as well as four 5-minute resting-state blood oxygen level-dependent scans, with their eyes open, fixed at a crosshair. Further details about the resting state imaging acquisition that varied by 3T scanner have been detailed previously (59)

#### Imaging Procedure: Processing

The ABCD DAIC performs centralized processing of MRI data from ABCD, using the Multi-Modal Processing Stream (61; see Supplement). After processing, between-network connectivity of the DMN and DAN was calculated by computing pairwise correlations between each ROI within the DMN and each ROI within the DAN, defined by the Gordon parcellation (62) These correlations were averaged and Fisher-Z transformed to generate a summary metric of DMN-DAN between-network connectivity strength.

**Figure 1.**
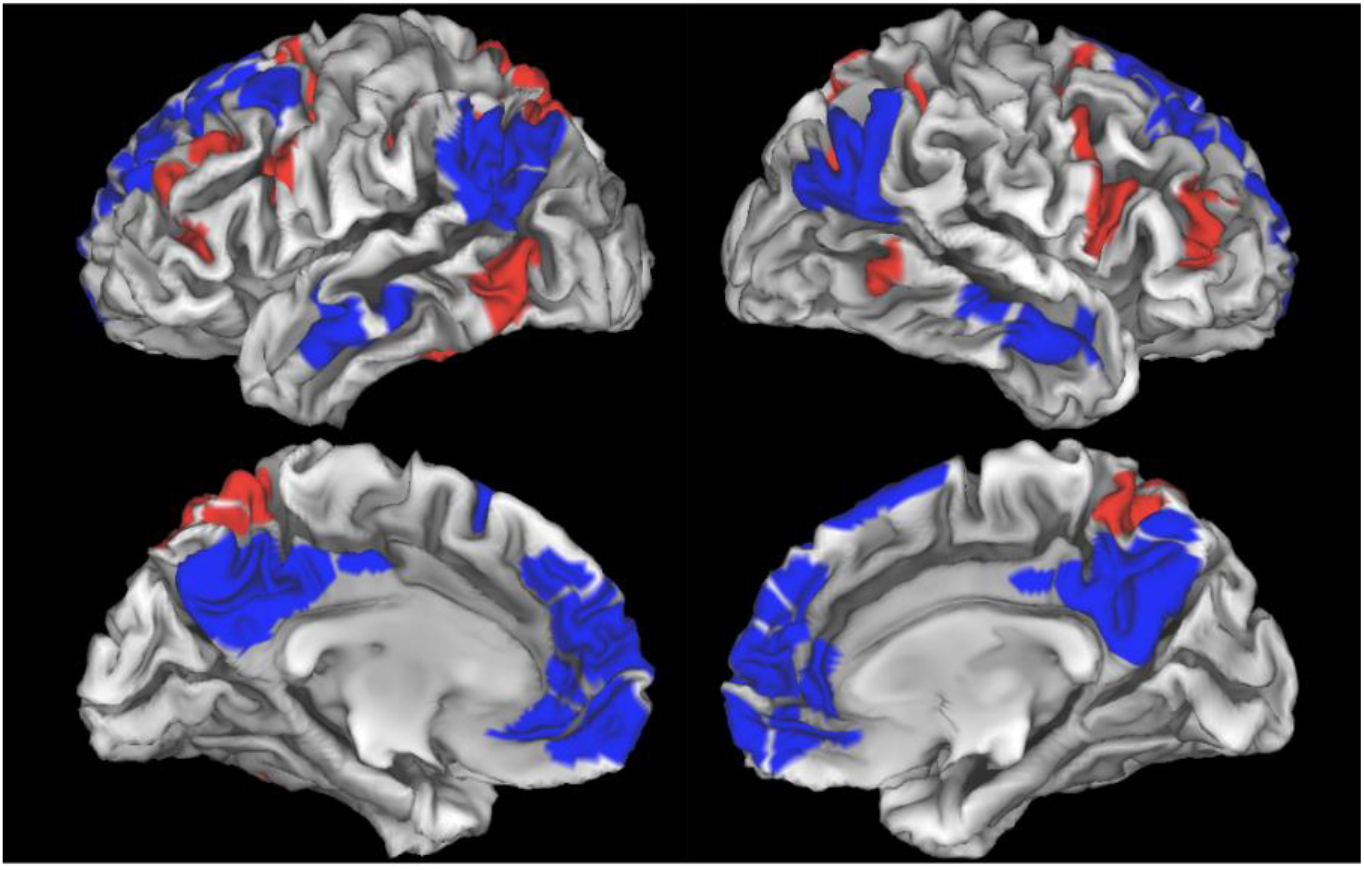
Default Mode Network (blue) and Dorsal Attention Network (red) from the Gordon Parcellation.

### Child Behavior Checklist: Attention and Externalizing Symptoms

Caregivers of the participants completed the Child Behavior Checklist (CBCL), a 112-item questionnaire used to detect emotional and behavioral problems in youth (63). Lastly, the longitudinal analyses use the CBCL attention and externalizing t-scores from follow-up years 1, 2, and 3. Demographic characteristics of participants by follow-up year, based on availability of the CBCL data, are included in the Supplement.

### Statistical Analyses

Linear mixed models were conducted in R version 4.1.1 lmer package (R Foundation for Statistical Computing, Vienna, Austria), with research site and family unit (nested within site, to account for 1336 twins and 23 triplets in this study sample) as random effects, and age, sex, race/ethnicity, parental income and parental education modeled as covariates (64). Imaging analyses additionally included mean framewise displacement (FD) as a covariate.

Linear mixed models were used to test the following cross-sectional predictions: (1a) across cognitive tasks (each in a separate model), lower IIV is associated the stronger DMN-DAN anti-correlation, and (1b) there would be an age effect, such that greater baseline age would be associated with stronger DMN-DAN anticorrelation and (1c) lower IIV. Furthermore, we tested the following hypotheses regarding associations between the lab-based measures and neurobehavioral symptoms: (2a) Greater IIV and/or (2b) weaker DMN-DAN anticorrelation are linked with worse attention symptoms at baseline. Lastly, we tested the predictions using longitudinal data: (3a) Across cognitive tasks (each in a separate model), greater IIV and (3b) weaker DMN-DAN anticorrelation, all assessed at baseline, will be associated with subsequent attention problems, 1-, 2-, and 3-years later, after accounting for baseline CBCL attention problems. Results are expressed as standardized coefficients with 95% confidence intervals. As a control analysis, we hypothesized that IIV and parent-reported attention problems would not be associated with between network connectivity between auditory and retrosplenial networks, since the Flanker, Dimensional Change Card Sort, and Pattern Comparison Processing speed tasks are not thought to engage either network (65).

## Results

### DMN-DAN Anticorrelation and Behavioral variability

As hypothesized, lower IIV across all three neurocognitive tasks, modeled in separate linear mixed models, was associated with stronger anticorrelation between the DMN and DAN (Table 2, see Supplement for estimated marginal means visualization). Within this model, older baseline age was associated with stronger DMN-DAN anticorrelation. Of the other covariates, head motion, sex, race/ethnicity, and family income were associated with DMN-DAN anticorrelation across models.

**Table 2.**
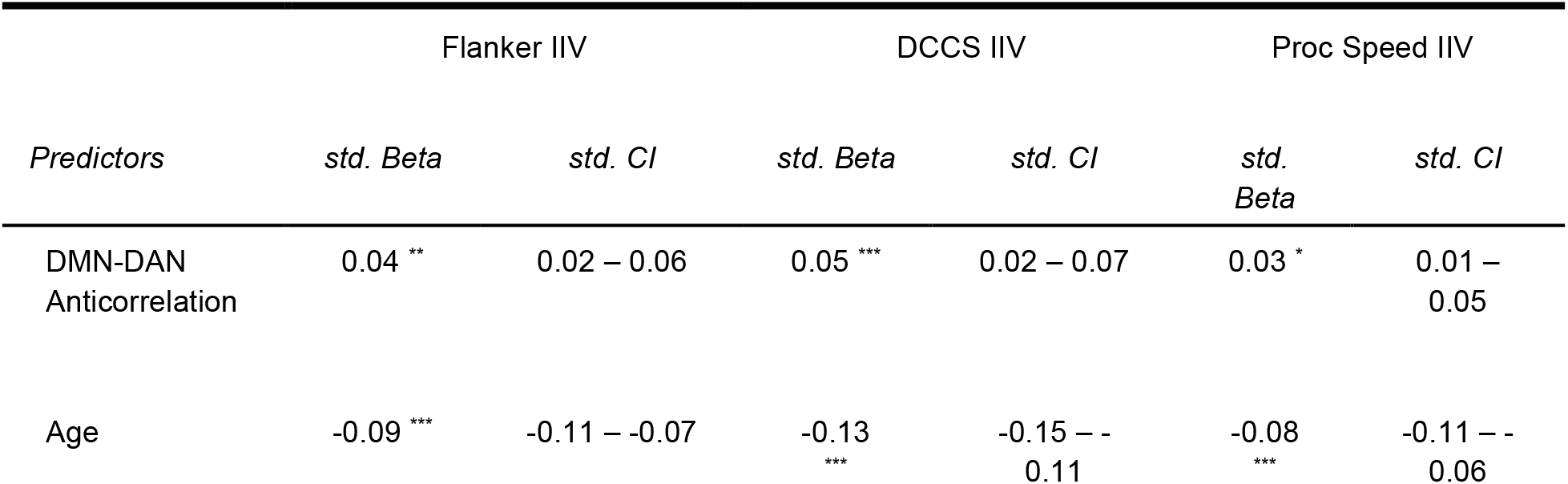
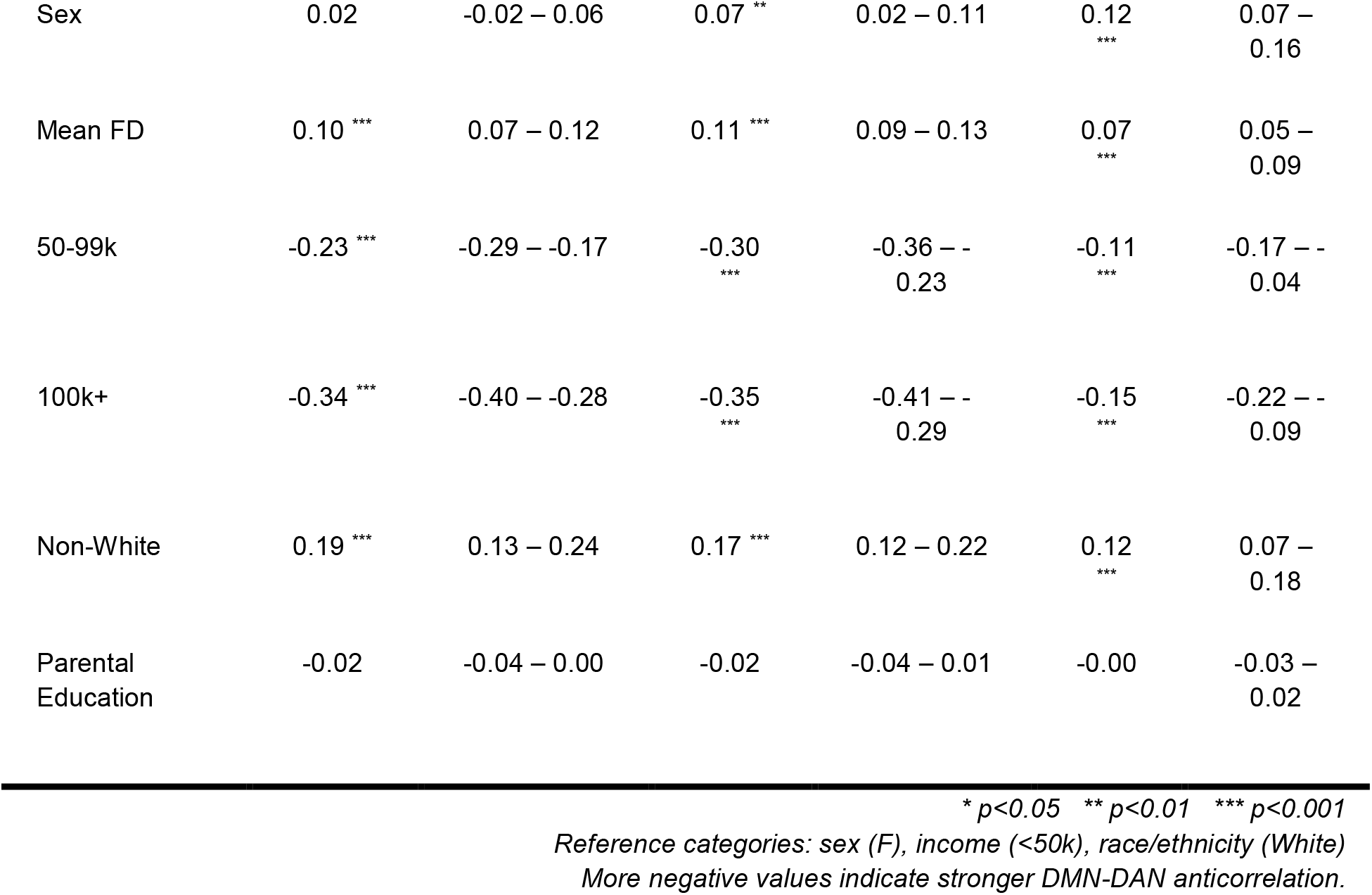
Associations between IIV and DMN-DAN Functional Connectivity.

| Predictors | Flanker IIV |  | DCCS IIV |  | Proc Speed IIV |  |
| --- | --- | --- | --- | --- | --- | --- |
|  | std. Beta | std. CI | std. Beta | std. CI | std. Beta | std. CI |
| DMN-DAN Anticorrelation | 0.04 ** | 0.02 – 0.06 | 0.05 *** | 0.02 – 0.07 | 0.03 * | 0.01 – 0.05 |
| Age | -0.09 *** | -0.11 – -0.07 | -0.13 *** | -0.15 – -0.11 | -0.08 *** | -0.11 – -0.06 |
| Sex | 0.02 | -0.02 – 0.06 | 0.07 ** | 0.02 – 0.11 | 0.12 *** | 0.07 – 0.16 |
| Mean FD | 0.10 *** | 0.07 – 0.12 | 0.11 *** | 0.09 – 0.13 | 0.07 *** | 0.05 – 0.09 |
| 50-99k | -0.23 *** | -0.29 – -0.17 | -0.30 *** | -0.36 – -0.23 | -0.11 *** | -0.17 – -0.04 |
| 100k+ | -0.34 *** | -0.40 – -0.28 | -0.35 *** | -0.41 – -0.29 | -0.15 *** | -0.22 – -0.09 |
| Non-White | 0.19 *** | 0.13 – 0.24 | 0.17 *** | 0.12 – 0.22 | 0.12 *** | 0.07 – 0.18 |
| Parental Education | -0.02 | -0.04 – 0.00 | -0.02 | -0.04 – 0.01 | -0.00 | -0.03 – 0.02 |

\* $p < 0.05$ \*\* $p < 0.01$ \*\*\* $p < 0.001$
Reference categories: sex (F), income (<50k), race/ethnicity (White)
More negative values indicate stronger DMN-DAN anticorrelation.

### Age and Behavioral Variability

We replicated neuropsychological literature showing that younger baseline age was also associated with greater IIV, across all three tasks after adjusting for covariates (4). Within these three models, there were also effects of race/ethnicity, parental income, and sex (Supplement). In a supplemental analysis, greater baseline age was also related to better accuracy (Supplement).

### Behavioral and Neural Associations with Attentional and Externalizing Symptoms at Baseline

Higher IIV across the Flanker, Processing Speed, and Dimensional Change Card Sort was associated with more severe baseline attention symptoms (Table 3). Furthermore, weaker DMN-DAN anticorrelation was associated with more severe attention symptoms (Table 3).

**Table 3.** Behavioral and Neural Associations with Attention Problems at Baseline.

| *Predictors* | CBCL Attention |  | CBCL Attention |  | CBCL Attention |  | CBCL Attention |  |
| --- | --- | --- | --- | --- | --- | --- | --- | --- |
|  | *std. Beta* | *std. CI* | *std. Beta* | *std. CI* | *std. Beta* | *std. CI* | *std. Beta* | *std. CI* |
| Flanker IIV | 0.12 *** | 0.10 –<br>0.14 |  |  |  |  |  |  |
| DCCS IIV |  |  | 0.13 *** | 0.11 –<br>0.15 |  |  |  |  |
| Proc Speed IIV |  |  |  |  | 0.11 *** | 0.09 –<br>0.13 |  |  |
| DMN-DAN<br>Anticorrelation |  |  |  |  |  |  | 0.07 *** | 0.05 –<br>0.09 |
| Mean FD |  |  |  |  |  |  | 0.05 *** | 0.02 –<br>0.07 |
| Age | 0.01 | -0.01 –<br>0.03 | 0.01 | -0.01 –<br>0.03 | 0.00 | -0.01 –<br>0.02 | 0.01 | -0.02 –<br>0.03 |
| Sex | 0.11 *** | 0.07 –<br>0.15 | 0.10 *** | 0.06 –<br>0.14 | 0.10 *** | 0.06 –<br>0.14 | 0.06 ** | 0.02 –<br>0.11 |
| 50-99k | -0.15 *** | -0.21 –<br>-0.10 | -0.14 *** | -0.20 –<br>-0.08 | -0.17 *** | -0.23 –<br>-0.12 | -0.17 *** | -0.23 –<br>-0.11 |
| 100k+ | -0.25 *** | -0.30 –<br>-0.19 | -0.24 *** | -0.30 –<br>-0.19 | -0.28 *** | -0.33 –<br>-0.22 | -0.28 *** | -0.34 –<br>-0.22 |
| Non-White | -0.01 | -0.05 –<br>0.04 | -0.01 | -0.06 –<br>0.03 | -0.00 | -0.05 –<br>0.04 | -0.01 | -0.06 –<br>0.04 |
| Parental<br>Education | -0.01 | -0.03 –<br>0.01 | -0.01 | -0.03 –<br>0.01 | -0.01 | -0.03 –<br>0.01 |  |  |

\* $p < 0.05$ \*\* $p < 0.01$ \*\*\* $p < 0.001$ Reference categories: sex (F), income (<50k), race/ethnicity (White) More negative values indicate stronger DMN-DAN anticorrelation.

**Table 4.** Prospective Behavioral Associations with Attentional symptoms, 1-3 years later.

|  | CBCL Attention Y1 |  | CBCL Attention Y2 |  | CBCL Attention Y3 |  |
| --- | --- | --- | --- | --- | --- | --- |
| *Predictors* | *std. Beta* | *std. CI* | *std. Beta* | *std. CI* | *std. Beta* | *std. CI* |
| (Intercept) | 0.05 * | 0.01 – 0.09 | 0.08 ** | 0.02 – 0.13 | 0.08 * | 0.01 – 0.15 |
| Flanker IIV | 0.02 ** | 0.01 – 0.03 | 0.03 ** | 0.01 – 0.04 | 0.03 * | 0.01 – 0.05 |
| Baseline CBCL<br>Attention | 0.73 *** | 0.71 – 0.74 | 0.68 *** | 0.66 – 0.70 | 0.61 *** | 0.59 – 0.63 |
| Age | -0.00 | -0.02 – 0.01 | 0.01 | -0.01 – 0.03 | 0.01 | -0.01 – 0.03 |
| Sex | 0.01 | -0.02 – 0.04 | -0.01 | -0.05 – 0.02 | -0.06 * | -0.10 – -<br>0.01 |
| 50-99k | -0.02 | -0.06 – 0.02 | -0.06 * | -0.10 – -<br>0.01 | -0.04 | -0.10 – 0.03 |
| 100k+ | -0.07 *** | -0.11 – -<br>0.04 | -0.06 * | -0.11 – -<br>0.01 | -0.06 * | -0.13 – -<br>0.00 |
| Non-White | -0.04 * | -0.07 – -<br>0.01 | -0.07 *** | -0.11 – -<br>0.03 | -0.05 | -0.10 – 0.00 |
| Parental Education | 0.00 | -0.01 – 0.02 | 0.00 | -0.02 – 0.02 | 0.01 | -0.01 – 0.04 |

\* $p < 0.05$ \*\* $p < 0.01$ \*\*\* $p < 0.001$
Reference categories: sex (F), income (<50k), race/ethnicity (White)
More negative values indicate stronger DMN-DAN anticorrelation.

In an exploratory analysis, higher IIV across all cognitive tasks and weaker (less negative) DMN-DAN anticorrelation was associated with more severe externalizing symptoms, all measured contemporaneously at baseline (Supplement). Of the covariates, several effects are consistent across all 8 models (Table 3 for Attentional Symptoms and Supplement for Externalizing Symptoms); sex, parental income, and race/ethnicity were associated with attention and externalizing symptoms.

### Prospective Behavioral and Neural Associations with Attentional Symptoms, 1-3 years later

Next, we evaluated if either IIV or DMN-DAN connectivity at baseline was predictive of attention symptoms, one, two, and three years after the baseline visit. For all three tasks, higher IIV at baseline predicted more severe attention symptoms at the participants’ 1-, 2-, and 3-year follow-up visits, after controlling for baseline attention problems (Flanker in Table 5, other tasks in Supplement). After accounting for baseline attention problems and covariates included in baseline models, DMN-DAN connectivity did not predict future attention symptoms (Supplement).

### Prospective Behavioral and Neural Associations with Externalizing Symptoms, 1-3 years later

Higher IIV at baseline was also associated with increased externalizing symptoms at the 1-year follow-up visit after controlling for baseline externalizing symptom scores, but not at later time points (Supplement). Furthermore, DMN-DAN anticorrelation did not predict future externalizing symptoms at any time point after controlling for baseline externalizing symptoms.

### Specificity and Robustness Analyses at Baseline

To demonstrate specificity, we tested an association between IIV and between-network connectivity between the auditory and retrosplenial networks. The cognitive tasks require visuospatial attention but not auditory processing. Furthermore, these tasks require in the moment attention, and not the range of cognitive functions linked with the retrosplenial network, including episodic memory, navigation, imagination and planning for the future (65). The auditory-retrosplenial network connectivity was not associated with Flanker IIV (p=0.753) or CBCL attention t-score (p=0.857) at baseline after accounting for the covariates used in previous models (Supplement).

To demonstrate the robustness of these results, we also applied a more stringent motion threshold which only included participants with 60% of resting-state frames or higher with framewise displacement less than 0.2 mm (n=7003). Greater DMN-DAN anticorrelation remained significantly associated with lower IIV in the Flanker and Dimensional Change Card Sort tasks, greater baseline age, and less severe baseline attention symptoms (Supplement).

## Discussion

The current study provides the first evidence linking anticorrelation between brain networks at rest and neurocognitive measures of attentional variability in early adolescence. Importantly, both lab-based resting state FC and neurocognitive markers of attentional variability were associated with parent-reported attention symptoms at baseline (ie. “fails to finish things they start”, “can’t concentrate”). We also provide novel evidence that worse scores on lab-based measures of attentional variability (IIV) at 9-10 years old, predict subsequent parent-reported attentional problems at 1, 2 and 3-year follow-up visits. Leveraging the ABCD cohort to evaluate longitudinal associations between FC, neurocognitive measures, and symptoms allows us to evaluate potential markers for attentional dysfunction in the general population and to understand the relationships between neural and behavioral development in adolescence.

To begin, we provide the first evidence linking stronger DMN-DAN anticorrelation and lower IIV in early adolescence, connecting resting-state and neurocognitive literature, which have been largely independent in studying attention variability. Previous studies have shown that the DMN and DAN networks have an intrinsically antiphase relationship, first established one year after birth, increasing during development, and decreasing in older adulthood (17, 22, 33-35, 66). Kelly et al. 2008 was the first to link stronger DMN-DAN anticorrelation with lower Flanker IIV in 20 adults. The current results extend this work to early adolescence and to other goal-directed, cognitive tasks, showing that stronger DMN-DAN anticorrelation is associated with greater baseline age and lower IIV across cognitive domains. Taken together, these findings suggest that segregation of the DMN and DAN may be relevant to cognitive development during early adolescence.

These results also indicate a remarkably consistent effect of age on neural and behavioral markers of attention, in a narrow age range from 9-11 years, that aligns with previous work in smaller samples (4-7). Neuropsychological task literature indicates that IIV decreases dramatically during childhood and adolescence, in conjunction with improved sustained attention (4-7). However, most of this literature focuses on IIV across the lifespan. Here, we focus on ages 9-11 and found that older baseline age was associated with lower IIV across the Flanker, Pattern Comparison Processing Speed, and Dimensional Change Card Sort Tasks. These consistent behavioral findings suggest that attentional variability is improving across different cognitive demands in early adolescence. Data on executive function task performance from more than 10,000 participants, aged 8-35, indicates that executive function most rapidly develops in early adolescence (46); our findings indicate robust age effects even within the narrow age band of 9-11.

The current study also found that higher IIV across three cognitive tasks is cross-sectionally associated with more severe attention symptoms. Previous work indicates that higher IIV robustly identifies ADHD vs. TD youth (2, 3), and as a result, takes a binary approach to attention problems (40, 47). However, continuous approaches are closer to the phenomena of attention and its underlying neurobiological processes (67-70). As such, this study takes a dimensional approach to include the full spectrum of attention problems in the ABCD sample (3, 71). Previous work in the ABCD study indicates that 949 individuals (7.99%) met ADHD diagnostic criteria, but the ABCD study also includes many youth with subclinical symptoms and youth with minimal attention problems (48). Despite these methodological differences, the current findings are consistent with previous work in children with ADHD and generalize to a representative sample of US youth (2).

The current results are the first to indicate that IIV assessed at age 9-10 is a consistent, proximal marker of worsening attention during early adolescence (1, 2, and 3 years later). These results align with two longitudinal, developmental studies (non-diverse, smaller samples) that controlled for attention symptoms at baseline and used IIV to predict overall functioning and symptoms of inattention (72,73). By controlling for attentional symptoms at baseline, the current results highlight the relationship between a lab-based measure of attentional variability in early adolescence and future worsening of attention symptoms, over and above early (parent-reported) attention deficits. In early adolescence, worsening attention problems may be a broad indicator of functional impairment, regardless of specific etiology or diagnoses that may emerge later on (8). Research on other neurodevelopmental disorders with onset in late adolescence (i.e, psychotic disorders) indicates that early intervention can be key for improving future symptom trajectories (74-79). Since ABCD will continue to collect data prospectively on participants through late adolescence, future work could examine IIV as a predictor of altered trajectories of cognitive development and mental health outcomes (8).

While we found that greater DMN-DAN anticorrelation was linked with lower contemporaneous attention problems, we did not find an association between DMN-DAN connectivity and future attention symptoms. The cross-sectional DMN-DAN FC -attention symptom association aligns with work using a previous release of the ABCD data (41). However, this work also stands in contrast with previous studies which have found no association between anticorrelated networks and attention symptoms, albeit in much smaller adult samples (80, 81). Using multivariate approaches may improve prediction abilities (82). Notably, Rosenberg et al. 2020 used a connectome-based predictive model to predict future sustained attention in adults (83). Therefore, other metrics of RSFC may be important neural predictors of symptom onset or progression.

In an exploratory analysis, we found that more severe externalizing symptoms were cross-sectionally associated with greater IIV and weaker DMN-DAN anticorrelation. Further, greater IIV at baseline predicted more severe externalizing symptoms at the following year, after controlling for externalizing symptoms at baseline. These results give credence to the possibility that attention is an important feature in self-regulation. Focusing on the task at hand, while suppressing internally-directed cognition, may be critical for controlling one’s behavior in relation to current and future goals (49-51).

### Limitations and Future Directions

This study leverages the largest imaging and behavior study of adolescence to date to examine a key developmental shift in attentional variability and its neural correlates; however, some limitations should be noted. Some argue that anticorrelation of default mode and task-positive networks may be a result of global signal regression (GSR) (84, 86). The current resting state literature highlights the importance of removing artifacts from motion, respiration, and physiological noise, which is particularly prominent in developmental populations (85). While GSR is a powerful tool to remove such spurious artifacts, it also can introduce negative correlations into the data, which has called into question whether DMN-DAN anticorrelation may be an artifact of GSR. However, anticorrelation between the default mode and task-positive networks has been detected in studies both with and without GSR, indicating that that anticorrelation is not a result of GSR (86-88). Because of the consistency of anticorrelation between default mode and task-positive networks, whether GSR is included or not, we have chosen to replicate and extend the majority of studies using GSR that show (i) development of anticorrelated networks or (ii) that anticorrelation is linked with superior cognitive performance (11, 33-35, 40, 47).

With regard to capturing attention problems, the CBCL inattention scale includes a variety of attention symptoms, including both hyperactivity and inattentiveness, which may warrant separate investigation in future studies (63). As a parent-report measure, the CBCL may be biased due to rater; multiple informants including teachers would be ideal to capture inattentive behavior across contexts.

The results of the current study demonstrate the functional significance of lab-based measures of attentional variability, linking them to attentional impairment in early adolescence. Increased intra-individual variability across three goal-directed behavioral tasks was robustly linked with more severe attention deficits at baseline as well as 1-, 2-, and 3-years later. While higher IIV is a well-known hallmark of current ADHD (2,3), these results show that IIV may be a useful index to identify worsening attention symptoms in the general population. Further, we have shown that stronger negative anticorrelation between the DMN and DAN is cross-sectionally associated with lower IIV across multiple tasks, better attention, and older age. These results suggest that the segregation of brain networks is associated with developmental improvements in cognition. In future studies, focusing on altered longitudinal patterns of DMN-DAN anticorrelation may be informative for predicting functional outcome.

## Acknowledgements and Disclosures

We thank Drs. Andrew Fuligni, Mirella Dapretto, and Adriana Galván for their feedback on these analyses and for their mentorship through the T32 for Brain and Behavioral Development during Adolescence. This work was supported by 1T32HD091059-01A1 (to SEC) and U01MH124639 (to CEB).

The authors report no biomedical financial interests or potential conflicts of interest.

