## Supplement for "Variability in cognitive task performance in early adolescence is associated with stronger between-network anticorrelation and future attention problems"

Manuscript

Click here to access/download;Supplement (Word doc);SChang_IIV_BiolPsych_Supp.docx

**Supplement Outline**

1. Note on missing ADHD diagnoses in the ABCD Data Release 4.0 (KSADS)
2. More detail on imaging acquisition/preprocessing done by ABCD
3. Demographics by site
4. Demographics by time point
5. Include winsorization details (how many outliers)
6. Analysis of IIV tasks - histograms, boxplot by site, accuracy, accuracy in relation to age
7. DMN-DAN connectivity - (a) by site, (b) by scanner type (c) mean FD by site (d) Aim 1 repeated analyses with more stringent motion threshold (e) Aim 2 repeated with stricter motion threshold
8. Aim 1: Estimated Marginal Means Visualization
9. Aim 1: Age effects on IIV
10. Aim 2: Neural and Behavioral Associations with Externalizing Symptoms
11. Linear Mixed Model Table - Predictive results for (a) DCCS and (b) Processing Speed tasks
12. Linear Mixed Model Table: Predictive results between IIV and CBCL Externalizing (Follow-up Year 1)
13. Linear Mixed Model Table - null results for DMN-DAN connectivity predicting future attention CBCL symptoms
14. Linear Mixed Model Table - specificity results; no association between correlation between auditory and retrosplenial networks and CBCL attention score
15. Details on neurocognitive tasks (Flanker, Dimensional Change Card Sort, Pattern Comparison Processing Speed)
16. Details on the Child Behavior Checklist

### 1. Missing ADHD diagnoses (KSADS)

Though ABCD used the Kiddie Schedule for Affective Disorders and Schizophrenia (KSADS) for the diagnostic criteria for psychiatric disorders, the current work used Data Release 4.0, in which these diagnoses were not available, due to several errors in the programming algorithm for calculating several psychiatric disorders, including ADHD. Regardless, our analyses focus on a dimensional approach toward attention dysfunction, so for this study we use the Child Behavior Checklist to capture attention problems on a continuous spectrum. Motion detection was conducted in real-time via the fMRI Integrated Real-time Motion Monitor (FIRMM), which allows study staff to adjust the scanning paradigm (more motion, the greater need for more data due to less usable data).

### 2. Supplementary Details on ABCD DAIC Preprocessing Stream

The scan parameters were optimized for consistency across all 19 study sites. T1-weighted imaging at 1mm^3 resolution was collected in the axial position at 256x256 matrix, 8 degree flip angle, and 2x parallel imaging. Other parameters varied by scanner type (Siemens: 176 slices, 256 × 256 FOV, 2500 ms TR, 2.88 ms TE, 1060 ms TI; Philips: 225 slices, 256 × 240 FOV, 6.31 ms TR, 2.9 ms TE, 1060 ms TI; GE: 208 slices, 256 × 256 FOV, 2500 ms TR, 2 ms TE, 1060 ms TI). Resting state fMRI scans were also collected in the axial position, at 2.4mm^3 resolution, 60 slices, 90x90 matrix, 216x216 FOV, 800ms TR, 30 ms TE, 52 degree flip angle, and 6-factor multiband acceleration.

Centralized data processing was carried out by the ABCD Data Analysis and Informatics Core. T1-weighted images underwent correction for gradient nonlinearity distortion and intensity inhomogeneity, and then were rigidly registered to a custom atlas. These images were then run through Freesurfer's automated pipeline to segment white matter, ventricle, and whole-brain ROIs. Resting state images were corrected for head motion, displacement, B0 distortions, and gradient non-linearity distortions, and then registered to the structural images. Initial frames were removed, and then the images were normalized by dividing the mean across time of each voxel. Signal from the estimated time courses (of six motion parameters, their derivatives and squares), quadratic trends, and the mean time course of white matter, ventricles, and whole brain were regressed out. Frames with displacement greater than 0.2mm were excluded. Temporal bandpass filtering (0.009-0.08Hz) was applied to these data.

These preprocessed time courses were sampled to FreeSurfer's cortical surface. Then, the Gordon parcellation's 13 functionally-defined networks were mapped, and the average time courses for Freesurfer's standard cortical and subcortical ROIs were calculated. Correlation values for each pair of ROIs were calculated, Fisher-Z transformed to z-statistics. Then for within network connectivity, the average correlation between each ROI pair within that network was calculated. Between network connectivity was calculated by the average correlation of pairs was calculated by including each ROI in one network with an ROI in the other network.

#### 3. CBCL/IIV/RSFC By Site

| Site Num | n | CBCL Attention t-score | CBCL Externalizing t-score | IIV (Flanker) | IIV (DCCS) | IIV (Processing Speed) | DMN DAN Correlation | Mean FD |
|---|---|---|---|---|---|---|---|---|
| site02 | 476 | 52.64(5.37) | 43.82(9.59) | 0.39(0.3) | 0.4(0.22) | 0.68(0.39) | -0.14(0.05) | 0.15(0.12) |
| site03 | 549 | 54.73(6.61) | 46.73(9.93) | 0.45(0.38) | 0.51(0.28) | 0.8(0.5) | -0.14(0.06) | 0.21(0.16) |
| site04 | 659 | 54.75(6.92) | 47.6(10.8) | 0.37(0.24) | 0.44(0.24) | 0.74(0.46) | -0.13(0.05) | 0.21(0.18) |
| site05 | 303 | 53.41(5.22) | 43.65(9.69) | 0.42(0.34) | 0.46(0.25) | 0.75(0.48) | -0.14(0.05) | 0.27(0.23) |
| site06 | 459 | 53.16(5.41) | 45.25(9.67) | 0.35(0.28) | 0.39(0.21) | 0.62(0.41) | -0.15(0.05) | 0.22(0.23) |
| site07 | 259 | 54.52(7.22) | 45.73(11.47) | 0.45(0.41) | 0.47(0.29) | 0.71(0.44) | -0.12(0.05) | 0.22(0.2) |
| site08 | 263 | 54.22(6.07) | 46.41(9.94) | 0.34(0.24) | 0.36(0.2) | 0.57(0.31) | -0.11(0.05) | 0.2(0.17) |
| site09 | 347 | 53.03(5.05) | 43.68(9.15) | 0.35(0.25) | 0.41(0.22) | 0.71(0.5) | -0.15(0.05) | 0.24(0.22) |
| site10 | 574 | 52.85(4.86) | 44.58(9.6) | 0.49(0.57) | 0.38(0.1) | 0.53(0.15) | -0.11(0.05) | 0.25(0.2) |
| site11 | 371 | 54.19(5.88) | 46.3(10.8) | 0.47(0.36) | 0.5(0.29) | 0.8(0.56) | -0.12(0.05) | 0.24(0.19) |
| site12 | 485 | 54.66(7.04) | 46.24(10.97) | 0.38(0.31) | 0.45(0.27) | 0.64(0.41) | -0.12(0.05) | 0.25(0.2) |

| | | | | | | | | |
|---|---|---|---|---|---|---|---|---|
| site13 | 624 | 53.29(5.54) | 44.58(9.83) | 0.4(0.32) | 0.42(0.25) | 0.67(0.42) | -0.12(0.05) | 0.22(0.18) |
| site14 | 462 | 52.37(4.41) | 43.32(9.16) | 0.32(0.25) | 0.37(0.2) | 0.6(0.37) | -0.15(0.05) | 0.22(0.18) |
| site15 | 313 | 55.3(7.83) | 48.75(12.09) | 0.6(0.5) | 0.6(0.35) | 0.98(0.7) | -0.11(0.05) | 0.31(0.27) |
| site16 | 936 | 54.15(6.55) | 47.44(9.77) | 0.34(0.23) | 0.39(0.21) | 0.62(0.39) | -0.16(0.05) | 0.15(0.13) |
| site18 | 290 | 53.33(5.18) | 44.84(9.7) | 0.41(0.35) | 0.38(0.18) | 0.62(0.37) | -0.11(0.05) | 0.21(0.18) |
| site20 | 562 | 53.31(5.56) | 44.96(10.58) | 0.39(0.27) | 0.46(0.27) | 0.71(0.41) | -0.14(0.05) | 0.25(0.23) |
| site21 | 483 | 53.27(5.76) | 44.36(10.33) | 0.43(0.35) | 0.42(0.24) | 0.74(0.52) | -0.15(0.05) | 0.27(0.24) |
| site22 | 29 | 54.1(4.83) | 47.24(8.45) | 0.22(0.2) | NaN(NA) | 0.36(0.15) | -0.1(0.05) | 0.17(0.1) |

### 4. Demographics By Year, based on available CBCL Attention t-score data

| | Year-1 Follow Up | Year-2 Follow-up | Year-3 Follow-up |
|---|---|---|---|
| Participants, N | 11207 | 8085 | 6133 |
| Sex, N(%) | | | |
| Female | 5344(47.68) | 3853(47.66) | 2899(47.27) |
| Male | 5863(52.32) | 4232(52.34) | 3234(53.73) |
| Baseline Age, N(%) | 9.91(62.58) | 9.94(62.15) | 9.97(61.94) |
| Race/Ethnicity, N(%) | | | |
| White | 5982(53.38) | 4508(55.76) | 3576(58.31) |
| Black | 1593(14.21) | 994(12.29) | 637(10.39) |
| Hispanic | 2216(19.77) | 1585(19.6) | 1175(19.16) |
| Asian | 241(2.15) | 178(2.2) | 136(2.2) |
| Other | 1173(10.47) | 820(10.14) | 609(9.93) |
| Parental Income, N(%) | | | |
| Less than $50k | 2923(26.08) | 2013(24.9) | 1428(23.28) |
| $50-99k | 2936(26.20) | 2192(27.11) | 1698(27.69) |
| $100k+ | 4433(39.56) | 3257(40.28) | 2593(42.28) |
| Refuse to report income | 462(4.12) | 325(4.02) | 221(3.6) |
| Don't know income | 452(4.03) | 298(3.69) | 193(3.15) |
| Education in years, N(SD) | 17.7(27.93) | 17.9(29.39) | 17.89(27.56) |
| CBCL Attention t-score, mean (SD) | 53.86(6.14) | 53.79(6.07) | 53.72(5.98) |
| CBCL Externalizing t-score, mean (SD) | 45.65(10.29) | 45.49(10.25) | 45.42(10.09) |
| Motion Inclusion, N(%) | 9264 (0.83) | 6714 (0.83) | 5099(0.83) |

### 5. Winsorization (how much of the data was replaced by 3SD)

| | Neurocognitive Task | % of Data Winsorized by 3SD |
|---|---|---|
| 1 | Flanker | 0.1987131 |
| 2 | Dimensional Change Card Sort | 2.2372680 |
| 3 | Processing Speed | 0.5680742 |

### 6. Neuropsychological Tasks - Supplementary plots and analyses

7a. Histograms

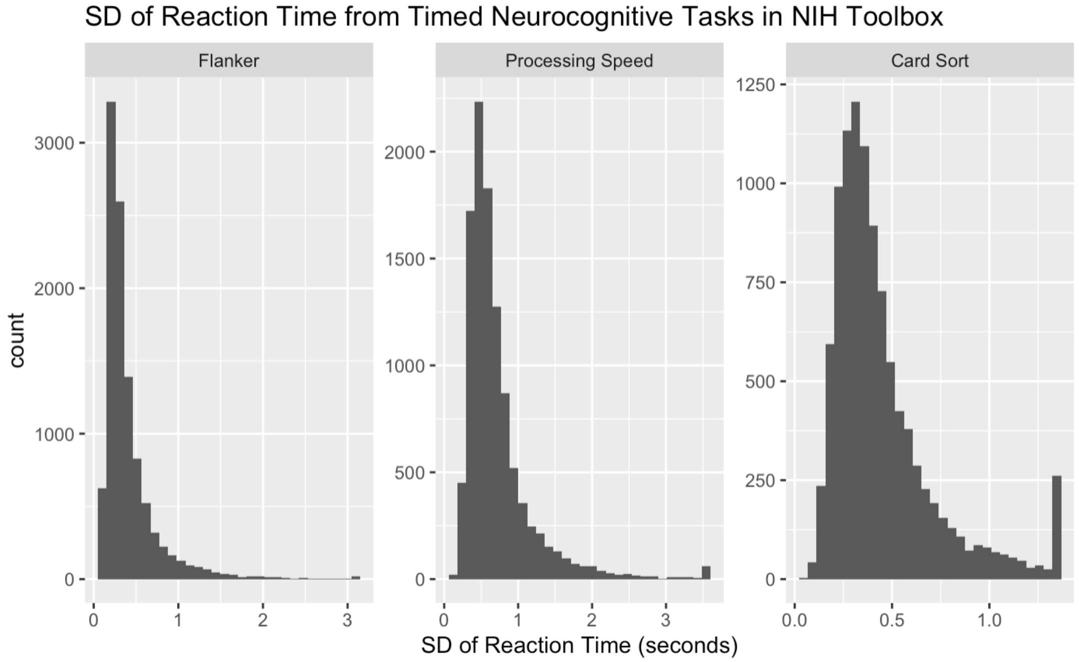


Notes: The standard deviations of reaction time across all correct trials were winsorized at 3 standard deviations to remove extremely large outliers.

**6b. IIV By Site Boxplots**

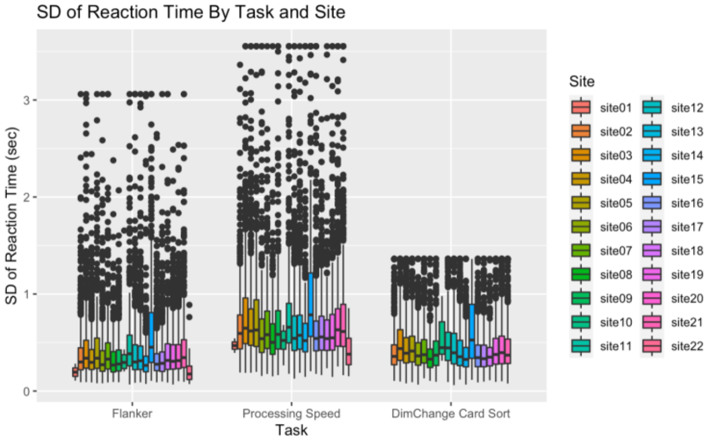


Note: Winsorized at 3SD, only correct trials included

**6c. Accuracy across all trials by task**

| Task | % of pts over 90% accurate |
|---|---|
| Flanker | 0.9847855 |
| Processing Speed | 0.9270519 |
| DimChange Card Sort | 0.8108740 |

**6d. Linear Mixed Models: Effect of Age on Accuracy**

| | Flanker Accuracy | | DCCS Accuracy | | Proc Speed Accuracy | |
|---|---|---|---|---|---|---|
| *Predictors* | *std. Beta* | *standardized CI* | *std. Beta* | *standardized CI* | *std. Beta* | *standardized CI* |
| (Intercept) | -0.04 | -0.11 – 0.02 | 0.02 | -0.06 – 0.10 | 0.12 *** | 0.06 – 0.19 |
| Age | 0.05 *** | 0.03 – 0.07 | 0.08 *** | 0.06 – 0.10 | 0.05 *** | 0.03 – 0.07 |
| Sex | -0.07 *** | -0.11 – -0.03 | -0.21 *** | -0.25 – -0.17 | -0.26 *** | -0.30 – -0.23 |
| Non-White | -0.09 *** | -0.14 – -0.05 | -0.13 *** | -0.17 – -0.08 | -0.00 | -0.05 – 0.04 |
| 50-99k | 0.13 *** | 0.08 – 0.19 | 0.18 *** | 0.13 – 0.24 | 0.02 | -0.04 – 0.07 |
| 100k+ | 0.18 *** | 0.13 – 0.24 | 0.23 *** | 0.18 – 0.29 | 0.04 | -0.02 – 0.09 |
| Parental Education | 0.01 | -0.01 – 0.03 | 0.02 * | 0.00 – 0.04 | -0.00 | -0.02 – 0.02 |

* *p<0.05* ** *p<0.01* *** *p<0.001*

*Reference categories: sex (F), income (<50k), race/ethnicity (White)*

### 7. DMN-DAN Correlation By Sites Included (no Phillips sites) -

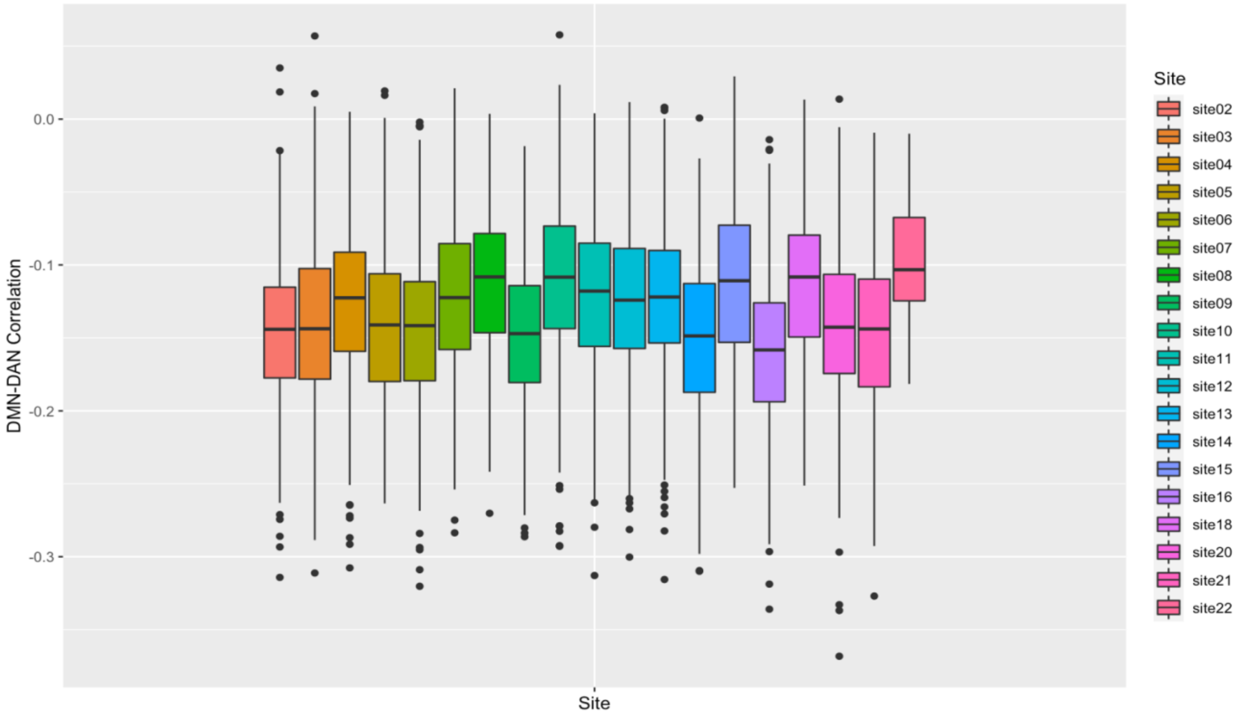


### 7b. DMN-DAN connectivity by scanner type

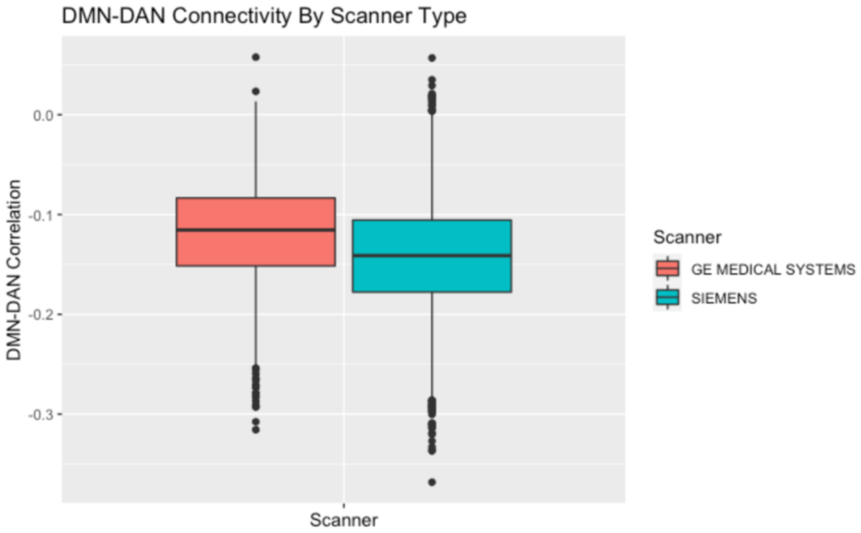

### 7c. Mean FD by Site

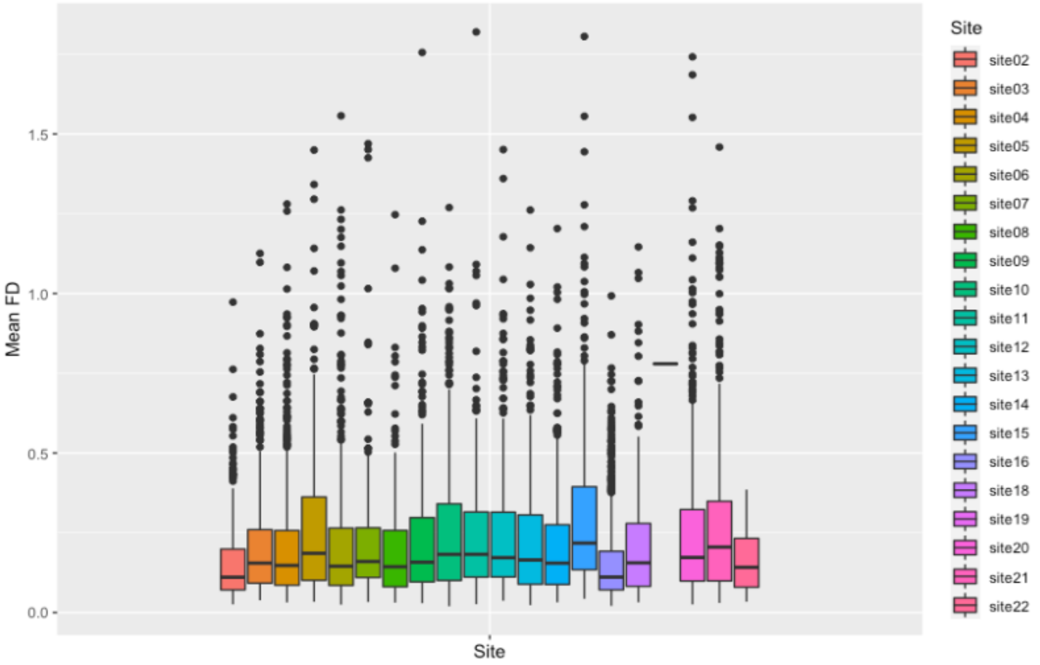

**7d. Aim 1a and 1b: Rerun with participants with at least 60% of frames with FD < 0.2mm (n=7003)**

**2. Stricter Motion Criteria (Aim 1)**

| | Flanker IIV | | DCCS IIV | | Proc Speed IIV | |
|---|---|---|---|---|---|---|
| *Predictors* | *std. Beta* | *standardized CI* | *std. Beta* | *standardized CI* | *std. Beta* | *standardized CI* |
| (Intercept) | 0.16 ** | 0.06 – 0.25 | 0.12 * | 0.02 – 0.22 | 0.02 | -0.09 – 0.12 |
| DMN-DAN Anticorrelation | 0.04 ** | 0.01 – 0.06 | 0.05 *** | 0.02 – 0.07 | 0.02 | -0.00 – 0.05 |
| Age | -0.10 *** | -0.12 – -0.07 | -0.13 *** | -0.16 – -0.11 | -0.08 *** | -0.11 – -0.06 |
| Sex | -0.00 | -0.05 – 0.05 | 0.05 * | 0.00 – 0.10 | 0.12 *** | 0.07 – 0.17 |
| Mean FD | 0.06 *** | 0.04 – 0.09 | 0.06 *** | 0.04 – 0.09 | 0.07 *** | 0.04 – 0.09 |
| 50-99k | -0.22 *** | -0.29 – -0.15 | -0.25 *** | -0.33 – -0.18 | -0.15 *** | -0.22 – -0.08 |
| 100k+ | -0.35 *** | -0.41 – -0.28 | -0.28 *** | -0.36 – -0.21 | -0.18 *** | -0.25 – -0.11 |
| Non-White | 0.18 *** | 0.12 – 0.24 | 0.15 *** | 0.09 – 0.20 | 0.11 *** | 0.05 – 0.16 |
| Parental Education | -0.02 | -0.05 – 0.00 | -0.06 *** | -0.09 – -0.03 | -0.01 | -0.03 – 0.02 |

** p<0.05 ** p<0.01 *** p<0.001*

*Reference categories: sex (F), income (<50k), race/ethnicity (White)*

*More negative values indicate stronger DMN-DAN anticorrelation.*

**7e. Aim 2b: Rerun with stricter motion threshold (participants with at least 60% of frames with FD < 0.2mm)**

| | CBCL Attention | | CBCL Externalizing | |
|---|---|---|---|---|
| *Predictors* | *std. Beta* | *standardized CI* | *std. Beta* | *standardized CI* |
| (Intercept) | 0.13 ** | 0.04 – 0.21 | 0.17 *** | 0.08 – 0.25 |
| DMN-DAN Anticorrelation | 0.07 *** | 0.04 – 0.09 | 0.04 ** | 0.01 – 0.07 |
| Age | 0.01 | -0.02 – 0.03 | -0.02 | -0.04 – 0.01 |
| Sex | 0.08 *** | 0.03 – 0.13 | 0.12 *** | 0.07 – 0.16 |
| Mean FD | 0.04 *** | 0.02 – 0.07 | 0.03 * | 0.00 – 0.05 |
| 50-99k | -0.19 *** | -0.26 – -0.13 | -0.22 *** | -0.29 – -0.16 |
| 100k+ | -0.30 *** | -0.36 – -0.23 | -0.37 *** | -0.43 – -0.30 |
| Non-White | 0.01 | -0.04 – 0.07 | -0.03 | -0.09 – 0.03 |
| Parental Education | -0.01 | -0.03 – 0.01 | -0.01 | -0.03 – 0.02 |

* *p<0.05* ** *p<0.01* *** *p<0.001*

*Reference categories: sex (F), income (<50k), race/ethnicity (White)*

*More negative values indicate stronger DMN-DAN anticorrelation.*

**8a. Aim 1: Estimated Marginal Means Visualization**

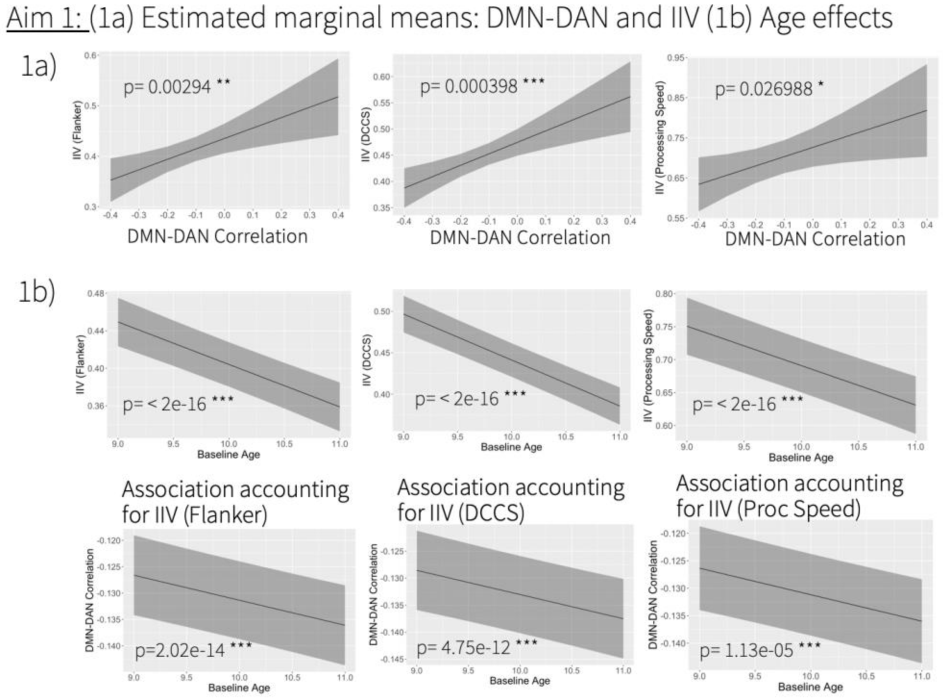


**8b. Aim 2: Estimated Marginal Means Visualization**

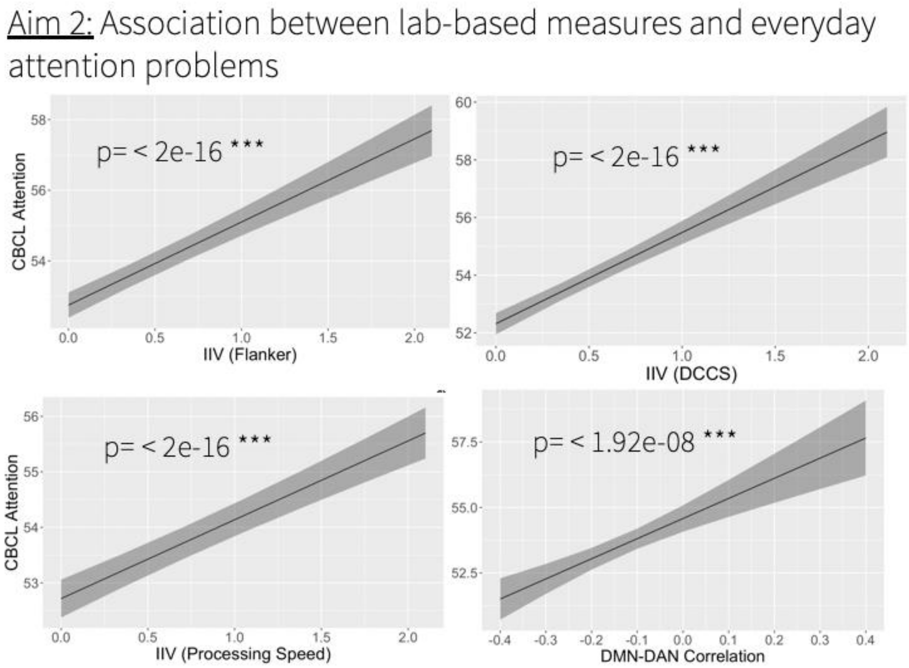

**8c. Aim 3: Estimated Marginal Means Visualization**

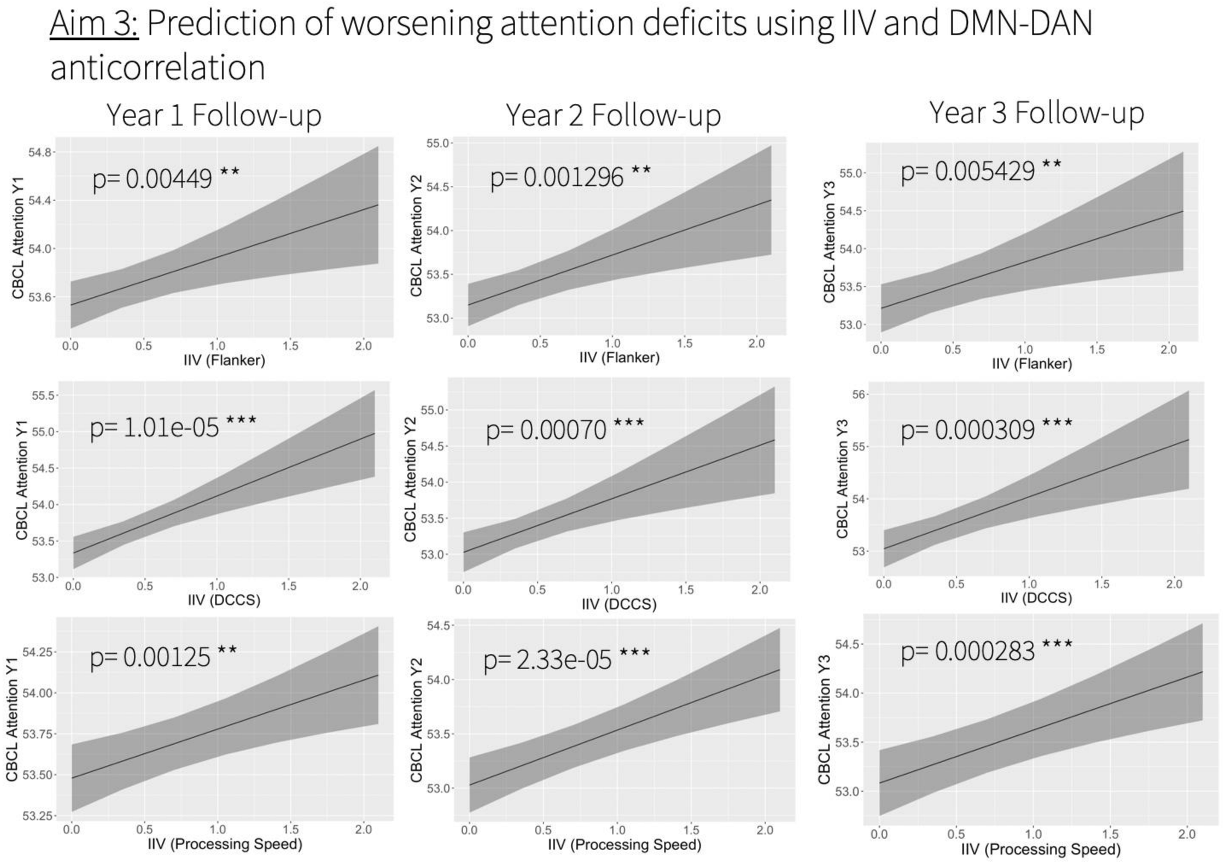

**9. Age effects on IIV performance**

| | DMN-DAN Anticorrelation | |
|---|---|---|
| *Predictors* | *std. Beta* | *standardized CI* |
| (Intercept) | -0.01 | -0.15 – 0.14 |
| Age | -0.06 *** | -0.08 – -0.04 |
| Sex | 0.22 *** | 0.18 – 0.26 |
| Mean FD | 0.19 *** | 0.17 – 0.21 |
| 50-99k | -0.06 | -0.11 – 0.00 |
| 100k+ | -0.13 *** | -0.18 – -0.07 |
| Non-White | 0.09 *** | 0.04 – 0.14 |
| Parental Education | -0.00 | -0.02 – 0.02 |

* $p<0.05$ ** $p<0.01$ *** $p<0.001$
*Reference categories: sex (F), income (<50k), race/ethnicity (White)*

**10. Behavioral and Neural Associations with Externalizing Symptoms at Baseline**

| | CBCL Externalizing | | CBCL Externalizing | | CBCL Externalizing | | CBCL Externalizing | |
|---|---|---|---|---|---|---|---|---|
| *Predictors* | *std. Beta* | *Std. CI* | *std. Beta* | *Std. CI* | *std. Beta* | *Std. CI* | *std. Beta* | *Std. CI* |
| (Intercept) | 0.17 *** | 0.10 – 0.24 | 0.18 *** | 0.11 – 0.25 | 0.18 *** | 0.11 – 0.25 | 0.17 *** | 0.09 – 0.26 |
| Flanker | 0.07 *** | 0.05 – 0.09 | | | | | | |
| DCCS | | | 0.10 *** | 0.08 – 0.12 | | | | |
| Proc Speed | | | | | 0.06 *** | 0.04 – 0.08 | | |
| DMN-DAN anticorrelation | | | | | | | 0.04 *** | 0.02 – 0.06 |
| Mean FD | | | | | | | 0.05 *** | 0.02 – 0.07 |
| Age | -0.02 | -0.04 – 0.00 | -0.01 | -0.03 – 0.01 | -0.02 * | -0.04 – -0.00 | -0.01 | -0.04 – 0.01 |
| Sex | 0.15 *** | 0.11 – 0.19 | 0.14 *** | 0.10 – 0.18 | 0.15 *** | 0.11 – 0.19 | 0.11 *** | 0.07 – 0.15 |
| 50-99k | -0.24 *** | -0.30 – -0.19 | -0.23 *** | -0.29 – -0.18 | -0.25 *** | -0.31 – -0.20 | -0.23 *** | -0.29 – -0.16 |
| 100k | -0.38 *** | -0.43 – -0.33 | -0.37 *** | -0.43 – -0.32 | -0.40 *** | -0.45 – -0.34 | -0.37 *** | -0.43 – -0.31 |
| Non-White | -0.06 ** | -0.11 – -0.01 | -0.08 ** | -0.12 – -0.03 | -0.06 * | -0.10 – -0.01 | -0.04 | -0.09 – 0.01 |

| | | | | | | |
|---|---|---|---|---|---|---|
| Parental Education | 0.00 | -0.02 – 0.02 | -0.01 | -0.03 – 0.01 | 0.00 | -0.02 – 0.02 |

** p<0.05   ** p<0.01   *** p<0.001*

*Reference categories: sex (F), income (<50k), race/ethnicity (White)*

*More negative values indicate stronger DMN-DAN anticorrelation.*

**11a. Attention Predictive Analysis: Dimensional Card Sort**

| | **CBCL Attention Y1** | | **CBCL Attention Y2** | | **CBCL Attention Y3** | |
|---|---|---|---|---|---|---|
| *Predictors* | *std. Beta* | *standardized CI* | *std. Beta* | *standardized CI* | *std. Beta* | *standardized CI* |
| (Intercept) | 0.05 ** | 0.01 – 0.09 | 0.07 * | 0.02 – 0.12 | 0.09 * | 0.02 – 0.16 |
| DCCS IIV | 0.03 *** | 0.02 – 0.05 | 0.03 ** | 0.01 – 0.05 | 0.04 *** | 0.02 – 0.06 |
| Baseline CBCL Attention | 0.73 *** | 0.71 – 0.74 | 0.68 *** | 0.66 – 0.69 | 0.60 *** | 0.58 – 0.63 |
| Age | -0.00 | -0.02 – 0.01 | 0.01 | -0.01 – 0.03 | 0.01 | -0.01 – 0.03 |
| Sex | 0.00 | -0.02 – 0.03 | -0.02 | -0.05 – 0.02 | -0.06 * | -0.10 – -0.01 |
| 50-99k | -0.02 | -0.06 – 0.02 | -0.05 | -0.10 – 0.00 | -0.05 | -0.11 – 0.02 |
| 100k+ | -0.07 *** | -0.11 – -0.03 | -0.05 * | -0.10 – -0.00 | -0.09 * | -0.16 – -0.02 |
| Non-White | -0.04 ** | -0.08 – -0.01 | -0.06 ** | -0.10 – -0.02 | -0.04 | -0.09 – 0.01 |
| Parental Education | 0.00 | -0.01 – 0.02 | 0.00 | -0.02 – 0.02 | 0.03 * | 0.00 – 0.05 |

* $p<0.05$   ** $p<0.01$   *** $p<0.001$

*Reference categories: sex (F), income (<50k), race/ethnicity (White)*

**11b. Attention Predictive Analysis: Processing Speed**

| | CBCL Attention Y1 | | CBCL Attention Y2 | | CBCL Attention Y3 | |
|---|---|---|---|---|---|---|
| *Predictors* | *std. Beta* | *standardized CI* | *std. Beta* | *standardized CI* | *std. Beta* | *standardized CI* |
| (Intercept) | 0.05 ** | 0.01 – 0.09 | 0.08 ** | 0.03 – 0.13 | 0.08 * | 0.02 – 0.15 |
| Proc Speed IIV | 0.02 ** | 0.01 – 0.04 | 0.04 *** | 0.02 – 0.05 | 0.04 *** | 0.02 – 0.06 |
| Baseline CBCL Attention | 0.73 *** | 0.71 – 0.74 | 0.68 *** | 0.66 – 0.69 | 0.61 *** | 0.59 – 0.63 |
| Age | -0.00 | -0.02 – 0.01 | 0.01 | -0.01 – 0.03 | 0.01 | -0.01 – 0.03 |
| Sex | 0.01 | -0.02 – 0.04 | -0.02 | -0.05 – 0.02 | -0.06 ** | -0.10 – -0.02 |
| 50-99k | -0.02 | -0.06 – 0.01 | -0.06 * | -0.11 – -0.01 | -0.04 | -0.10 – 0.03 |
| 100k+ | -0.08 *** | -0.12 – -0.04 | -0.07 ** | -0.11 – -0.02 | -0.07 * | -0.13 – -0.00 |
| Non-White | -0.04 * | -0.07 – -0.01 | -0.07 *** | -0.11 – -0.03 | -0.05 * | -0.10 – -0.00 |
| Parental Education | 0.00 | -0.01 – 0.02 | -0.00 | -0.02 – 0.02 | 0.01 | -0.01 – 0.04 |

* *p<0.05* ** *p<0.01* *** *p<0.001*

*Reference categories: sex (F), income (<50k), race/ethnicity (White)*

*More negative values indicate stronger DMN-DAN anticorrelation.*

**12. Association between IIV and CBCL Externalizing (Follow-up Year 1)**

| | CBCL Externalizing | | CBCL Externalizing | | CBCL Externalizing | |
|---|---|---|---|---|---|---|
| *Predictors* | *std. Beta* | *standardized CI* | *std. Beta* | *standardized CI* | *std. Beta* | *standardized CI* |
| (Intercept) | 0.06 * | 0.01 – 0.10 | 0.05 * | 0.01 – 0.10 | 0.06 ** | 0.02 – 0.10 |
| Flanker | 0.02 * | 0.00 – 0.03 | | | | |
| DCCS | | | 0.02 * | 0.00 – 0.03 | | |
| Proc Speed | | | | | 0.02 * | 0.00 – 0.03 |
| Baseline CBCL Externalizing | 0.73 *** | 0.71 – 0.74 | 0.73 *** | 0.71 – 0.74 | 0.73 *** | 0.71 – 0.74 |
| Age | 0.01 | -0.00 – 0.03 | 0.01 | -0.00 – 0.03 | 0.01 | -0.00 – 0.02 |
| Sex | 0.01 | -0.02 – 0.03 | 0.01 | -0.02 – 0.03 | 0.00 | -0.02 – 0.03 |
| 50-99k | -0.05 * | -0.09 – -0.01 | -0.04 * | -0.08 – -0.00 | -0.05 ** | -0.09 – -0.01 |
| 100k+ | -0.10 *** | -0.14 – -0.06 | -0.10 *** | -0.14 – -0.06 | -0.11 *** | -0.14 – -0.07 |
| Non-White | -0.01 | -0.04 – 0.02 | -0.01 | -0.04 – 0.03 | -0.01 | -0.04 – 0.03 |
| Parental Education | -0.00 | -0.01 – 0.01 | 0.01 | -0.01 – 0.02 | -0.00 | -0.01 – 0.01 |

** p<0.05   ** p<0.01   *** p<0.001*
*Reference categories: sex (F), income (<50k), race/ethnicity (White)*
*More negative values indicate stronger DMN-DAN anticorrelation.*

**13. Null results: DMN-DAN Correlation association with future CBCL attention symptoms**

| | **CBCL Attention Y1** | | **CBCL Attention Y2** | | **CBCL Attention Y3** | |
|---|---|---|---|---|---|---|
| *Predictors* | *std. Beta* | *standardized CI* | *std. Beta* | *standardized CI* | *std. Beta* | *standardized CI* |
| (Intercept) | 0.03 | -0.01 – 0.08 | 0.06 * | 0.01 – 0.12 | 0.09 * | 0.01 – 0.17 |
| DMN-DAN Anticorrelation | 0.01 | -0.01 – 0.03 | 0.00 | -0.02 – 0.02 | 0.01 | -0.02 – 0.04 |
| CBCL Baseline Attention | 0.73 *** | 0.71 – 0.75 | 0.68 *** | 0.66 – 0.70 | 0.60 *** | 0.57 – 0.62 |
| Age | 0.00 | -0.01 – 0.02 | 0.01 | -0.01 – 0.03 | 0.02 | -0.00 – 0.05 |
| Sex | 0.00 | -0.03 – 0.04 | -0.02 | -0.06 – 0.02 | -0.09 *** | -0.14 – -0.05 |
| Mean FD | 0.01 | -0.01 – 0.03 | 0.01 | -0.01 – 0.03 | 0.04 *** | 0.02 – 0.07 |
| Non-White | -0.03 | -0.07 – 0.00 | -0.05 * | -0.10 – -0.01 | -0.05 | -0.10 – 0.01 |
| 50-99k | -0.00 | -0.05 – 0.04 | -0.03 | -0.09 – 0.02 | -0.03 | -0.09 – 0.04 |
| 100k+ | -0.05 * | -0.09 – -0.01 | -0.06 * | -0.12 – -0.01 | -0.06 | -0.12 – 0.01 |

| | | | | | | |
|---|---|---|---|---|---|---|
| Parental Education | 0.00 | -0.01 – 0.02 | -0.00 | -0.02 – 0.02 | 0.02 | -0.01 – 0.04 |

* p<0.05 ** p<0.01 *** p<0.001

*Reference categories: sex (F), income (<50k), race/ethnicity (White)*

*More negative values indicate stronger DMN-DAN anticorrelation.*

**14. Specificity Analysis - No association between auditory-retrosplential between network FC and IIV**

| | Flanker | | CBCL Attention | |
|---|---|---|---|---|
| *Predictors* | *std. Beta* | *standardized CI* | *std. Beta* | *standardized CI* |
| (Intercept) | 0.14 ** | 0.05 – 0.23 | 0.12 ** | 0.05 – 0.20 |
| Auditory-Retrosplenial FC | 0.01 | -0.01 – 0.03 | -0.00 | -0.03 – 0.02 |
| Age | -0.09 *** | -0.12 – -0.07 | 0.00 | -0.02 – 0.02 |
| Sex | 0.03 | -0.02 – 0.07 | 0.08 *** | 0.03 – 0.12 |
| Mean FD | 0.10 *** | 0.08 – 0.13 | 0.06 *** | 0.04 – 0.08 |
| Non-White | 0.19 *** | 0.14 – 0.24 | -0.01 | -0.06 – 0.04 |
| 50-99k | -0.23 *** | -0.30 – -0.17 | -0.18 *** | -0.24 – -0.12 |
| 100k+ | -0.34 *** | -0.41 – -0.28 | -0.29 *** | -0.35 – -0.23 |
| Parental Education | -0.02 | -0.04 – 0.00 | -0.01 | -0.03 – 0.01 |

* p<0.05 ** p<0.01 *** p<0.001

*Reference categories: sex (F), income (<50k), race/ethnicity (White)*

**15. Further Detail on Neurocognitive Tasks**

For ABCD, these tasks were specifically designed and validated to be used across childhood and adolescence and were administered via iPad, in English with trained study staff.

| Task Image | Task Description |
|---|---|
| 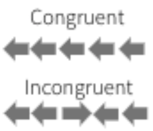  | The Flanker Task measures attention/cognitive control (Eriksen and Eriksen 1974; Fan et al. 2002; Rueda et al. 2004). On each trial, five arrows appear in a row. Four flanking stimuli (two on the outer left, two on the outer right) are facing in the same direction, either left or right. Participants indicate by touch input if the middle arrow is facing either left or right, regardless of the direction of the flanking arrows. Scoring is based on both speed and accuracy. |
| 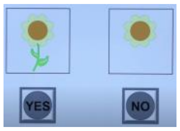  | The Pattern Comparison Processing Speed Task (Carlozzi et al. 2015; Salthouse et al. 1991) measures rapid visual processing: Participants are shown two pictures and asked to determine by touch input whether the two pictures are identical or not. The goal is to complete the task as quickly and as accurately as possible. |
| 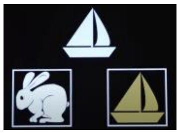 | The Dimensional Change Card Sort task measures cognitive flexibility (Zelazo 2006). During each trial, two different objects of two different colors appear at the bottom of the screen. Participants sort a third object in the middle of the screen by either color or shape to match one of the other two objects. The task consists of two initial blocks, sorting on one dimension for each block, and then a final block consists of a set of trials with alternating sorting dimensions.The score is based on both accuracy and reaction time. |

**16. Details on the Child Behavior Checklist**

Each item on the checklist was rated from 0 (not true) to 2 (very true) based on the last 6 months. Based on summing the scores from relevant items, the CBCL includes subscales for attention problems, rule-breaking behavior, aggressive behavior, anxious/depressed, withdrawn/depressed, somatic complaints, social problems, and thought problems. For this study, ourwe primary analysesily used the CBCL attention subscale, which includes items like "Can't concentrate, can't pay attention for long" and "Fails to finish things they start". In an exploratory analysis, we also examined the CBCL externalizing subscale, the sum of rule-breaking and aggressive behavior items, in relation to IIV. For these analyses, we used the t-scores for attention and externalizing subscales; all CBCL subscale t-scores have a mean of 50 and standard deviation of 10 (Achenbach 1507).
